# Modulation of genetic- and toxicant-induced Parkinson’s disease outcomes by SV2C in *C. elegans*

**DOI:** 10.64898/2026.11.20.767914

**Authors:** Ava C. Knickerbocker, Haejung Chung, Gary W. Miller, Meghan L. Bucher

**Author notes:** Corresponding author: Columbia University, 650 W 168th St, BB-1602, New York, NY 10032, United States.

## Abstract

Several proteins involved in dopaminergic neurotransmission have been implicated in Parkinson’s disease (PD), including synaptic vesicle glycoprotein 2C (SV2C). At least seven SNPs in SV2C are associated with PD, including variants that modify: PD risk, PD risk in GBA carriers, PD severity, contribution of environmental factors (e.g., nicotine) to PD, and PD patient response to L-DOPA and dopamine therapies. Although the specific function of SV2C remains unknown, there is significant evidence supporting a role of SV2C in mediating vesicular dopamine content, vesicular trafficking, and neurotransmitter release. Furthermore, SV2C directly associates with α-synuclein and mediates the content of neurotoxicants within synaptic vesicles. Taken together, these data suggest that SV2C plays a critical role in mediating PD risk and pathogenesis. To investigate whether SV2C mediates PD outcomes, transgenic *Caenorhabditis elegans* (*C. elegans*) expressing human SV2C, human α-synuclein, or both, were exposed to the PD-associated neurotoxicants 1-methyl-4-phenylpyridinium (MPP^+^), rotenone, and paraquat, and evaluated for PD outcomes including lethality and dopamine-mediated motor behavior. Although SV2C-expressing *C. elegans* showed a shorter lifespan at baseline, SV2C expression conferred organism-level protection against MPP^+^ and paraquat-induced lethality. While α-synuclein expression exacerbated lethality following MPP^+^ and paraquat exposure, SV2C provided partial rescue of lethality in α-synuclein-expressing *C. elegans*. However, no differences in lethality were observed based on genotype following rotenone exposure. Interestingly, sublethal toxicant exposure did not reveal a protective effect of SV2C expression on motor behavior for any of the toxicants. To determine whether SV2C’s modulation of vesicular uptake mediated the protection against lethality from PD-associated neurotoxicants observed in *C. elegans, in vitro* assays of vesicular dopamine uptake dynamics were performed. SV2C expression resulted in a protective shift in toxicant-induced uptake inhibition following MPP^+^ and rotenone exposure; however, there was no paraquat-induced inhibition of vesicular uptake observed *in vitro*. Taken together, these data suggest that while SV2C plays a protective role in mediating organism-level survival in *C. elegans* following MPP^+^ and paraquat exposure, the mechanisms of protection and the role of SV2C in modulating vesicular dopamine dynamics and dopamine-mediated behaviors following toxicant exposure require further analysis. Future studies can evaluate additional PD outcomes in these exposure models and investigate SV2C-mediated mechanisms of protection such as regulation of neurotransmitter handling and oxidative stress.

## Introduction

Parkinson’s disease (PD) is a progressive neurodegenerative disease characterized by motor deficits resulting from degeneration of dopaminergic neurons within the nigrostriatal pathway. Development of PD from solely genetic factors is rare; approximately 90% of cases are sporadic, believed to be due to aging, environmental factors, and gene-environment interactions.^1,2^ Several toxicants associated with PD have been well characterized in preclinical experimental models, including 1-methyl-4-phenylpyridinium (MPP^+^), rotenone, and paraquat. These compounds trigger neuronal dysfunction and degeneration through a variety of mechanisms, including mitochondrial dysfunction and oxidative stress, and result in the aberrant formation of α-synuclein.^3–6^

The damage resulting from toxicant exposures can be compounded upon by dysregulated dopamine, which has the potential to act as an endogenous neurotoxin when it accumulates in the cytosol.^7– 10^ In healthy conditions, the cytosolic pool of dopamine is minimized by sequestration and retention within synaptic vesicles, processes mediated by vesicular monoamine transporter 2 (VMAT2) and synaptic vesicle glycoprotein 2C (SV2C), respectively.^11,12^ Several SNPs in VMAT2 and SV2C have been associated with PD, and there is significant evidence of cytosolic dopamine accumulation in PD.^13–24^ It is believed that cytosolic dopamine contributes to neuronal dysfunction and degeneration due to degradation and enzymatic- and auto-oxidation, processes that generate reactive oxygen species and reactive metabolites.^8,25,26^ Furthermore, cytosolic dopamine contributes to the formation of aberrant α-synuclein, suggesting that regulation of dopamine is critical to maintaining neuronal health in PD.^25,27–29^

Deficits in VMAT2-mediated dopamine sequestration result in increased dopamine turnover, motor deficits, neurodegeneration, and the induction of PD-associated pathogenic mechanisms of degeneration.^25,30–32^ However, increased VMAT2 expression is neuroprotective against MPTP and methamphetamine.^33,34^ Deficits in SV2C result in reduced vesicular dopamine uptake, reduced vesicular dopamine retention, reduced evoked dopamine release, and increased vulnerability to MPTP.^11,35^ SV2C also co-immunoprecipitates with α-synuclein, and multimeric α-synuclein is present in the midbrain of SV2C knockout mice, suggesting SV2C influence α-synuclein aggregation.^35^ Taken together, these data support the hypothesis that the dysregulation of dopamine sequestration contributes to PD pathogenesis and progression, and that VMAT2 and SV2C may mediate neurotoxicity from toxicant exposures.

Understanding the relationship between gene variants and environmental exposures associated with PD will provide key insight into PD pathogenesis, progression, treatment, and prevention. To that end, this study investigated the ability of SV2C expression to mediate PD outcomes in *Caenorhabditis elegans* (*C. elegans*) models, including dopaminergic α-synuclein expression, and following exposure to MPP^+^, paraquat, and rotenone. After the generation of novel *C. elegans* and *in vitro* models, organism lethality and DA-mediated behaviors were evaluated to test whether SV2C confers protection against MPP^+^, paraquat, and rotenone exposure. Additionally, *in vitro* assays were performed for mechanistic insight to determine whether SV2C protects against toxicant-induced impairments in vesicular uptake dynamics. This study provides foundational evidence that SV2C expression in *C. elegans* provides organism-level protection against toxicant exposure, with future studies providing additional insight in the mechanisms of this protection.

## Methods and Materials

### *C. elegans* strains and husbandry

The N2 and BZ555 (*egIs1* [Pdat-1::GFP]) strains were obtained from the Caenorhabditis Genetics Center, which is funded by the NIH Office of Research Infrastructure Programs (P40 OD010440). The UA44 (*baIn11* [Pdat-1::alpha-syn + Pdat-1::gfp]) strain was provided as a gift from Dr. Catharine Rankin’s laboratory at the University of British Columbia. The SV2C (JBRA; Pdat-1::hSC2V::mScarlet::dat-1u) expressing strain was derived by InVivo Biosystems, and the SV2C::BZ555 and SV2C::UA44 crosses were generated in the Miller Laboratory. Throughout the manuscript, the strains are referred to as follows: BZ555 is GFP, UA44 is GFP::α-syn, SV2C::BZ555 is GFP::SV2C, and SV2C::UA44 is GFP::α-syn::SV2C. All *C. elegans* strains were maintained on agar plates with normal growth medium (NGM) at 20°C with OP50 *E. coli* as a food source.

### Genotyping

Individual *C. elegans* were analyzed for genotype by lysis with the Extract-N-Amp™ Tissue PCR Kit (Sigma-Aldrich, XNAT2). Each *C. elegans* was incubated in Extraction Solution and Tissue Preparation Solution at 55°C for 10 min, followed by 95°C for 3 min. Lysis was neutralized with Neutralization Solution B, and the resulting lysate was used directly as a PCR template. PCR was performed using OneTaq® 2X Master Mix with Standard Buffer (New England Biolabs, M0482S). The primers for SV2C (forward: GACTTACAACGGAGAGGCTAAC; reverse: CCTCCGATCATCCAGAACATAC) yielded a PCR product of 732 base pairs, and the primers for α-synuclein (forward: AGAAGCAGCAGGAAAGACAA; reverse: GTTCGTAGTCTTGATACCCTTCC) yielded a PCR product of 332 base pairs. Amplification was performed with an initial denaturation at 94°C for 2 min, followed by 30 cycles of 94°C for 15 sec, 58°C for 15 sec, and 68°C for 45 sec, with a final extension at 68°C for 5 min. PCR products were resolved on 1% E-Gel™ EX agarose gels (Invitrogen) using a 1-kb Plus DNA Ladder (Thermo Fisher Scientific).

### Synchronization of the population

Bleach synchronization was performed as previously described.^36^ Briefly, gravid adult *C. elegans* were collected with sterilized water, centrifuged to remove the supernatant, and treated with bleach solution (250 mM KOH and 0.5% hypochlorite prepared in sterile water) until the *C. elegans* cuticle dissolved. Tubes were centrifuged and the bleach solution was removed without disturbing the egg pellet. The egg pellet was washed with sterile water three times before a final resuspension in M9 solution (22 mM KH2PO4, 42 mM Na2HPO4, 85 mM NaCl, and 1 mM MgSO4 prepared in sterile water) and transfer to 50 mL conical tubes. Conical tubes were kept on a 20°C shaking incubator overnight for eggs to hatch to larval stage 1 (L1).

### Body length analysis

Body length analysis was performed as previously described.^36^ Briefly, synchronized populations of *C. elegans* were placed on agar pads on microscope slides while immobilized with 5 mM levamisole (MP Biomedicals). Brightfield images were acquired of *C. elegans* using an APX100 microscope (Evident) at 4x. L1s were analyzed from bleach synchronization and developmental arrestment, the combined larval stage 2 (L2) and larval stage 3 (L3) was analyzed at 24 h post-L1 plating, the larval stage 4 (L4) was analyzed at 48h post-L1 plating, and the Day One Adult stage was analyzed at 72 h post-L1 plating. Body length was determined using ImageJ with Fiji and the segmented line tool, where a line was drawn through the midline of each *C. elegans* from head to tail. Data was analyzed in GraphPad Prism 11 using two-way ANOVA with Tukey’s post hoc multiple comparison’s test.

### Eggs *in utero* analysis

Fecundity was assessed as previously described.^36^ Briefly, individual gravid *C. elegans* were placed in one well of a clear U-shaped 96-well plate containing 20% bleach solution prepared in sterile water. Once the *C. elegans* cuticle dissolved, the number of eggs in the well were counted using an EVOS*fl* microscope at 4x and 10x. The Day One Adult stage was analyzed at 72 h post-L1 plating, Day Two Adult stage was analyzed at 96 h post-L1 plating, and the Day Three Adult stage was analyzed at 120 h post-L1 plating. Data was analyzed in GraphPad Prism 11 using two-way ANOVA with Tukey’s post hoc multiple comparison’s test.

### Lifespan analysis

Lifespan analysis was adapted from Kalia *et al*. 2022.^37^ Briefly, approximately 50 L4 *C. elegans* per strain were placed on a 6cm NGM plate spotted with OP50. The experiment was performed in triplicate, with three plates each containing 50 *C. elegans* per strain. Plates were evaluated every 24 h to count the number of alive and dead *C. elegans* on each plate. *C. elegans* were transferred to new plates daily for the first six days of adulthood to isolate the original population from future generations. Post-reproductive stage *C. elegans* were then transferred to new plates twice a week to decrease the likelihood of contamination and replenish food. *C. elegans* were evaluated as dead if they did not respond to gentle touch with a platinum wire and were censored from analysis if they were missing from the plate, lost during transfer, showed internal hatching of L1s, or died of a non-age-related cause (e.g. rupturing at the vulva). Data analysis was performed in GraphPad Prism 11 using Kaplan-Meier survival analysis and Holm-Šídák’s multiple comparisons test.

### Toxicant exposures in *C. elegans*

Synchronized L1 *C. elegans* were grown on NGM plates for 48 h until the L4 stage. Liquid culture plates were prepared using S-complete buffer, 5-fluorodeoxyuridine (FUdR), ampicillin, and UV inactivated NEC937 with water or DMSO as solvent controls, 5 mM, 7.5 mM, and 10 mM MPP^+^, 15 mM, 20 mM, and 25 mM paraquat, or 20 µM, 30 µM, 40 µM, and 50 µM rotenone. Approximately 50 L4 *C. elegans* were added to each well of the liquid culture plate, and the plate was maintained at 20°C on a shaking incubator for 48 h.

### Lethality assays

*C. elegans* exposed to toxicants for 48 h were collected onto 6 cm unspotted NGM agar plates and liquid was left to evaporate. Once evaporated, *C. elegans* were counted and evaluated as dead if they did not respond to gentle touch with a platinum wire. Data was analyzed in GraphPad Prism 11 using two-way ANOVA with Tukey’s post hoc multiple comparison’s test.

### WormLab behavior

WormLab behavioral analysis was performed to evaluate overall motor behavior and dopamine-mediated behaviors. Following sublethal toxicant exposure in liquid culture, *C. elegans* were transferred to a NGM agar plate for behavioral analysis. Videos were taken for 30 s at a frame rate of 7 frames per second, and videos were analyzed using WormLab (MBF Bioscience) software. Data was analyzed in GraphPad Prism 11 using two-way ANOVA with Tukey’s post hoc multiple comparison’s test.

### Cell culture

Human Embryonic Kidney cells (HEK293) were generated by GenScript by stable transfection of a vector expressing human Green Fluorescent Protein (GFP)-tagged Dopamine Transporter (DAT) and human Vesicular Monoamine Transporter 2 (VMAT2) under Puromycin selection. An additional stable cell line was generated on this background to include expression of human Synaptic Vesicle Glycoprotein 2C (SV2C) under Geneticin selection. Cells were maintained in an incubator at 37°C with 5% CO2 on poly-D-lysine coated 10 cm cell culture plates in media consisting of Dulbecco’s Modified Eagle Medium (DMEM) with 4.5 g/L glucose (Corning), 10% fetal bovine serum (Fisher), .5% penicillin– streptomycin (Sigma), and selection antibiotics.

### Western blot

HEK-DAT-VMAT2 and HEK-DAT-VMAT2-SV2C cells were washed with ice-cold PBS and lysed in M-PER lysis buffer supplemented with protease inhibitors. Cell lysates were sonicated and centrifuged at 14,000 rpm for 15 m at 4°C, and the supernatants were collected for protein analysis. Protein samples were loaded onto NuPAGE 4–12% Bis-Tris gels. Lysates were mixed with NuPAGE LDS Sample Buffer (4X; Invitrogen, Cat. No. NP0007) and NuPAGE Sample Reducing Agent (10X; Invitrogen, Cat. No. NP0009) and separated by SDS-PAGE using NuPAGE 4–12% Bis-Tris precast gels (Thermo Fisher Scientific) with NuPAGE MOPS SDS Running Buffer at 200 V for 30 m. Following separation, proteins were transferred onto nitrocellulose membranes (Bio-Rad) using NuPAGE Transfer Buffer (20X; Invitrogen, Cat. No. NP0006) supplemented with 10% methanol at 17 V for 1 h. Membranes were blocked with 5% milk for 1 h and incubated overnight with primary antibodies against VMAT2 (OriGene, TA500506; 1:1,000), SV2C (Proteintech, 21271-1-AP; 1:1,000), GFP (Invitrogen, A11122), and β-actin (cell signaling 8457; 1:3000), followed by IRDye 680RD- and IRDye 800CW-conjugated secondary antibodies (LI-COR; 1:15,000) for 1h at room temperature. Blots were imaged using a LI-COR imaging system.

### FFN206 uptake assays

Assays of vesicular uptake were performed as previously described.^11,38^ HEK-DAT-VMAT2 or HEK-DAT-VMAT2-SV2C cells were seeded on black-walled half-area 96-well plates (Corning) at 50,000 cells per well in culture media 24 h before assay. For the uptake assay, culture media was removed, and cells were incubated with solvent control (DMSO or sterile water) or tetrabenazine (Sigma), rotenone (Sigma), paraquat (Sigma), and 1-methyl-4-phenylpyridinium (MPP^+^) (Sigma) at a range of doses (0.001 µM - 100 µM) prepared in DMEM with 4.5 g/L glucose, sodium pyruvate, without L-glutamine, and phenol red (Corning) for 30 m. Following 30 m pre-treatment, FFN206 (Tocris) was added to a final concentration of 1.33 µM for 1 h. At the end of the 1 h co-incubation period, the media containing the treatment and FFN206 was removed and replaced with media containing 1:25 trypan blue. Plates were scanned using a BioTek Synergy H1 multiplate reader at 37°C using end-point fluorescent intensity measurements with excitation λ = 369 and emission λ = 464. Fluorescent values underwent background subtraction and normalization to solvent control. Normalized data underwent IC_50_ calculation performed in GraphPad Prism 11 by nonlinear regression ([inhibitor] vs. normalized response), and regressions were compared to determine significant differences in IC_50_ value or Curve of Best Fit. Data points were excluded only in cases of pipetting errors or cell detachment during assay, as confirmed by microscopy.

### Statistics

Graphing and statistical analyses were performed using GraphPad Prism 11.

## Results

A transgenic strain of *C. elegans* expressing human SV2C was created to investigate the role of SV2C in mediating genetic- and toxicant-induced Parkinson’s disease outcomes in *C. elegans. C. elegans* do not endogenously express SV2 proteins; however, the *C. elegans* genome includes a closely related orthologous protein, SVOP.^39^ The transgenic strain of *C. elegans* expressing SV2C was generated by InVivo Biosystems, with SV2C expression under the dopamine transporter (DAT) promoter to restrict expression to the dopaminergic neurons, and a transgene marker appearing as green fluorescent protein (GFP) in the pharynx (Figure 1A). Additionally, an mScarlet tag was included on SV2C for fluorescent detection and localization (Figure 1A). Western blot analysis was performed to confirm the presence of SV2C in the transgenic strain (Figure 1B).

**Figure 1.**
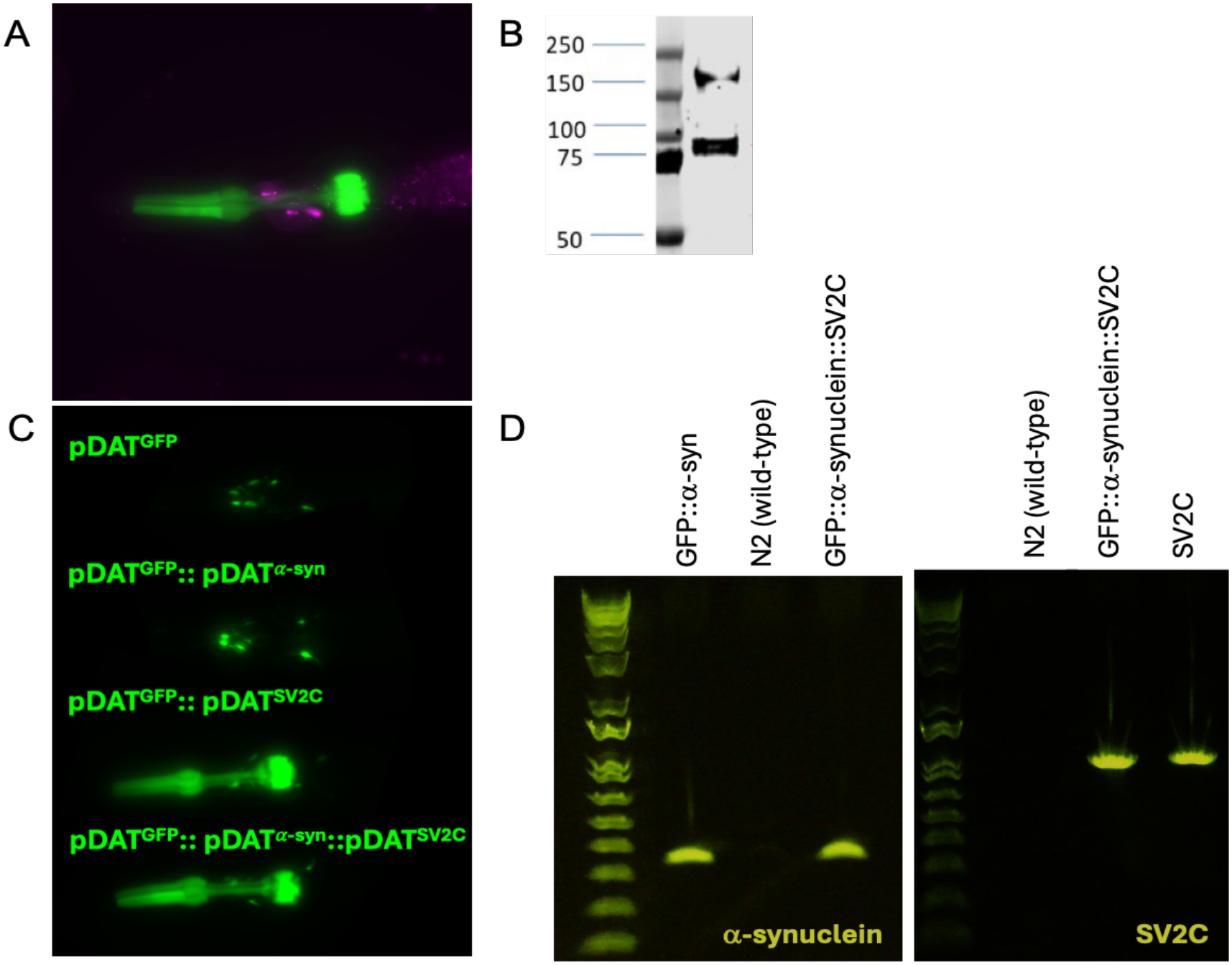
Validation of transgenic SV2C-expressing *C. elegans* strains. A. 60x microscopy image of mScarlet-tagged SV2C in JBRA (pDAT^SV2C^) strain with GFP transgene marker in the pharynx. B. Western blot of SV2C expression in JBRA (pDAT^SV2C^) strain. C. 20x microscopy images of GFP in dopamine neurons and transgene marker in the pharynx in strains expressing GFP, SV2C, and α-synuclein. D. Genotype confirmation of α-synuclein and SV2C expression with negative (N2 wild-type) and positive (UA44, JBRA) controls.

Following confirmation of SV2C expression in the transgenic strain, male *C. elegans* were generated by heat shock to breed additional SV2C-expressing strains. Breeding plates were prepared with male transgenic SV2C-expressing *C. elegans and* hermaphrodite BZ555 (*egIs1* [Pdat-1::GFP]) or UA44 (*baIn11* [Pdat-1::alpha-syn + Pdat-1::gfp]) *C. elegans* to generate a control strain with both SV2C expression and a fluorescent dopaminergic neuron marker, or an experimental strain with both SV2C and α-synuclein expression in addition to the fluorescent dopaminergic neuron marker, respectively. Successful integration of the genotype was confirmed by observing the GFP pharynx transgene marker, indicating SV2C expression, and GFP expression within the dopamine neurons, indicating successful integration with the BZ555 or UA44 strains. Individual *C. elegans* with both fluorescent markers were selectively passaged, followed by additional single-offspring passages for two generations until a homologous population resulting from an individual *C. elegans* was generated with 100% fluorescent marker presentation (Figure 1C). Genotype confirmation on these strains validated the successful integration of SV2C and α-synuclein in the intended strains (Figure 1D).

To determine whether the integration of SV2C and α-synuclein into the *C. elegans* genome impacted morphological features or reproductive behaviors, the strains were evaluated for body length across developmental stages, the number of eggs *in utero* across three days of adulthood, and lifespan. There were no morphological defects observed in the SV2C and α-synuclein expressing strains, such as vulval defects, dumpy body shape, or roller phenotype (data not shown). Brightfield microscopy analysis of body length demonstrated no significant differences in body length from larval stage L1 through larval stage L4 (Figure 2A). At Day One Adulthood, the BZ555 (pDAT^GFP^; GFP) strain had significantly longer body length compared to UA44 (pDAT^GFP^::pDAT^α-syn^; ; GFP::α-syn) (p < 0.0001), SV2C (pDAT^GFP^::pDAT^SV2C^; GFP::SV2C) (p < 0.0001), and SV2C::UA44 (pDAT^GFP^::pDAT^SV2C^::pDAT^α-syn^; GFP::α-syn::SV2C) (p < 0.0001). Additionally, the GFP::α-syn strain had significantly longer body length compared to SV2C (pDAT^GFP^::pDAT^SV2C^) (p < 0.05) (Two-way ANOVA, Tukey’s post hoc multiple comparisons test). However, the magnitude of body length difference observed corresponded to less than 10% difference in length for all strains, suggesting that these differences in body length are not biologically relevant.

**Figure 2.**
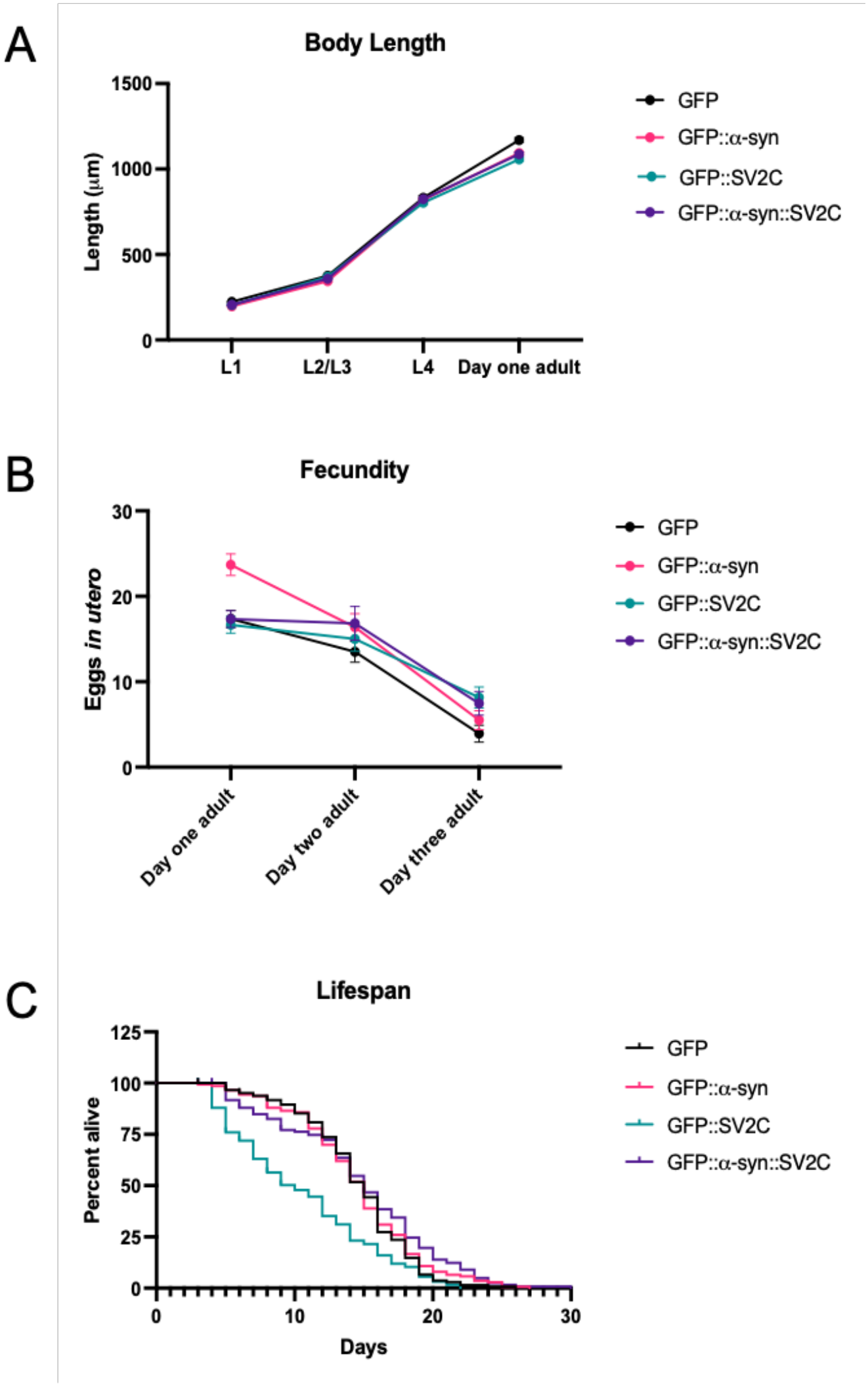
Morphological and biological characterization of transgenic SV2C expressing *C. elegans* strains. A. Body length (µm) measured at developmental stages L1, L2/L3, L4, and at day one of gravid adulthood. Each data point represents the mean and standard error of the mean compiled across three experimental replicates totaling between n = 48 - 109 *C. elegans* per strain at each developmental stage. Two-way ANOVA with Tukey’s post hoc multiple comparisons test. B. Fecundity assay measured by the number of eggs *in utero* at day one, day two, and day three of gravid adulthood. Each data point represents the mean and standard error of the mean compiled across three experimental replicates totaling n = 30 *C. elegans* per strain at each age. Two-way ANOVA with Tukey’s post hoc multiple comparisons test. C. Survival plot demonstrating lifespan as a percent of the number of *C. elegans* alive across days of adulthood. Each line represents compiled data from three experimental replicates, each beginning with 45-51 *C. elegans*. Mantel-Cox test with Holm-Šídák’s multiple comparisons test.

While observation of the GFP::α-syn::SV2C strain revealed a higher proportion of naturally occurring male offspring (data not shown), a fecundity assay to evaluate egg laying revealed no significant differences in the number of eggs *in utero* for hermaphrodite GFP::α-syn::SV2C *C. elegans* through three days of adulthood (Figure 2B). However, at Day One Adulthood, the GFP::α-syn strain showed significantly more eggs *in utero* compared to GFP (p < 0.01), GFP::SV2C (p < 0.001), and GFP::α-syn::SV2C (p < 0.01) equivalent to roughly 6-7 more eggs *in utero* than other genotypes (Two-way ANOVA, Tukey’s post hoc multiple comparisons test).

Lastly, analysis of lifespan demonstrated a significantly shorter lifespan for the GFP::SV2C strain compared to all other genotypes (Mantel-Cox with Holm-Šídák’s multiple comparisons test, p < 0.0001) (Figure 2C). The GFP::SV2C strain reached 0% alive by 22 days of adulthood compared to 26 days of adulthood for GFP, 27 days of adulthood for GFP::α-syn, and 30 days of adulthood for GFP::α-syn::SV2C. Despite the observed increase in eggs *in utero* in the GFP::α-syn strain, and the decreased lifespan observed for the GFP::SV2C strain, the GFP::α-syn::SV2C strain was not significantly morphologically, biologically, or reproductively different from the control GFP strain, suggesting a bi-directional rescue in the observed biological effects from combined gene expression.

After confirming that the SV2C and α-synuclein expressing *C. elegans* were morphologically unaffected, lethality was evaluated following toxicant exposure to determine whether SV2C mediates organism-level survival. After an initial dose response for MPP^+^, paraquat, and rotenone was performed, three doses were selected for lethality analysis. *C. elegans* were exposed to the compounds at the L4 stage and the number of dead *C. elegans* was evaluated for each dose following 48h of exposure. The GFP::α-syn strain showed increased vulnerability to MPP^+^ compared to the GFP::SV2C strain at the 7.5mM (p < 0.05) and 10mM (p < 0.01) doses (Two-way ANOVA with Tukey’s post hoc multiple comparisons test; Figure 3A). These data suggest that α-synuclein expression increases vulnerability to MPP^+^ exposure, and SV2C expression protects against MPP^+^ exposure.

**Figure 3.**
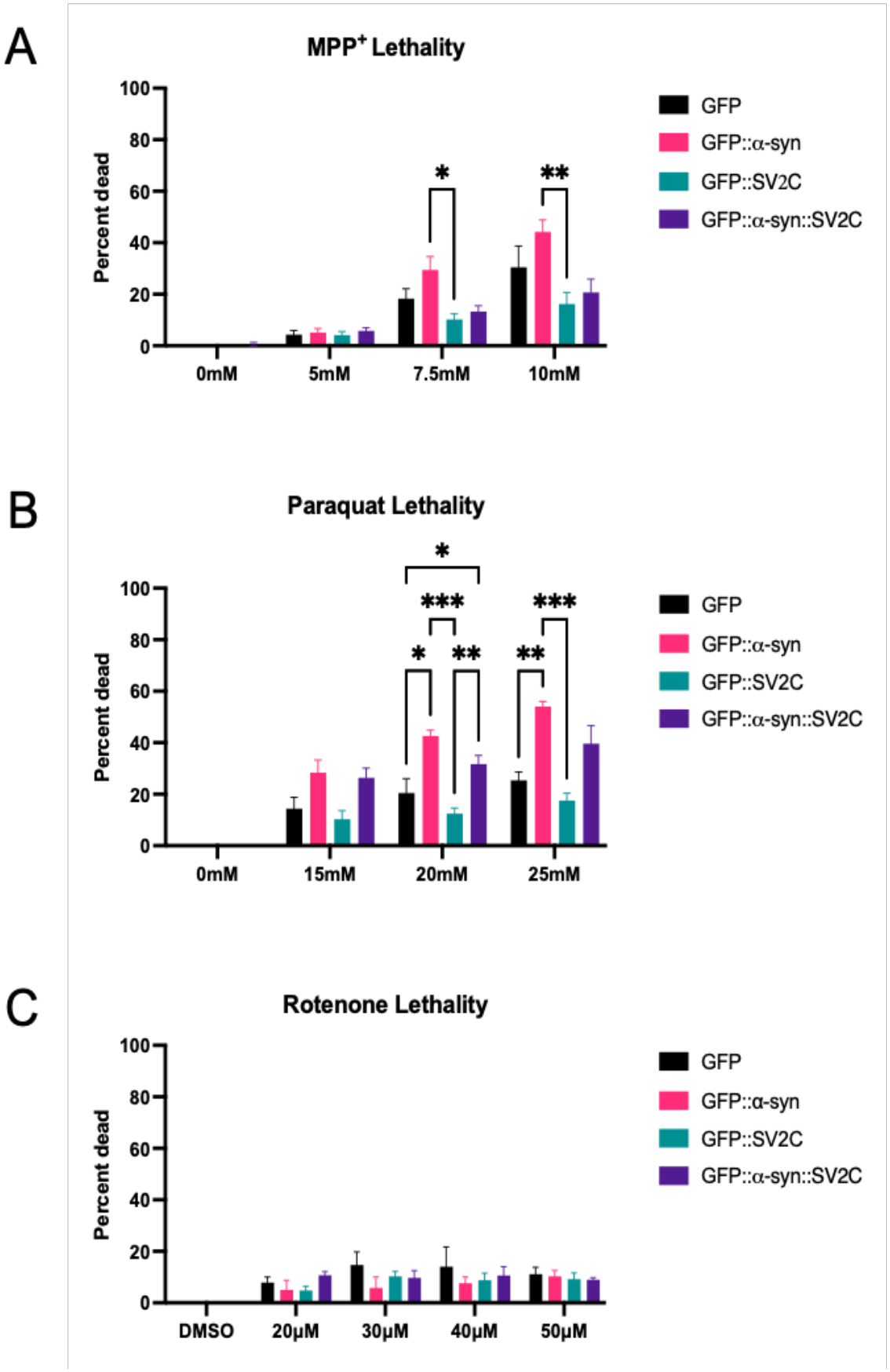
Lethality following MPP^+^, paraquat, and rotenone exposure. A. Lethality represented as the percent of *C. elegans* dead following 48h exposure to 5mM, 7.5mM, and 10mM MPP^+^. Data compiled across five experimental replicates with n = 18 - 40 *C. elegans* per group in each experiment. Data represented as the mean and standard error of the mean. Two-way ANOVA with Tukey’s post hoc multiple comparisons test, *p < 0.05, **p < 0.01. B. Lethality represented as the percent of *C. elegans* dead following 48h exposure to 15mM, 20mM, and 25mM paraquat. Data compiled across five experimental replicates with n = 24 - 56 *C. elegans* per group in each experiment. Data represented as the mean and standard error of the mean. Two-way ANOVA with Tukey’s post hoc multiple comparisons test, *p < 0.05, **p < 0.01, ***p < 0.001. C. Lethality represented as the percent of *C. elegans* dead following 48h exposure to 20µM, 30µM, 40µM, and 50µM rotenone. Data compiled across five experimental replicates with n = 23 - 58 *C. elegans* per group in each experiment. Data represented as the mean and standard error of the mean. Two-way ANOVA with Tukey’s post hoc multiple comparisons test, n.s.

Similarly, the GFP::α-syn strain showed increased vulnerability to paraquat compared to the GFP strain at the 20mM (p < 0.05) and 25mM (p < 0.01) doses and compared to the SV2C (pDAT^GFP^::pDAT^SV2C^) strain at the 20mM (p < 0.001) and 25mM (p < 0.001) doses (Figure 3B). Additionally, the GFP::α-syn::SV2C strain showed increased vulnerability to paraquat exposure compared to the GFP strain at the 20mM dose (p < 0.05) and compared to the GFP::SV2C strain at 20mM (p < 0.01) (Two-way ANOVA with Tukey’s post hoc multiple comparisons test; Figure 3B). Taken together, these data suggest that α-synuclein expression increases vulnerability to paraquat exposure, and SV2C expression protects against paraquat exposure

For all of the doses of rotenone tested, there were no differences in vulnerability observed between all of the *C. elegans* strains tested (n.s.) (Two-way ANOVA with Tukey’s post hoc multiple comparisons test; Figure 3C). Additionally, the strains demonstrated lower and more variable vulnerability to the rotenone exposure compared to MPP^+^ and paraquat. Despite the low vulnerability to rotenone exposure observed, rotenone was included in additional analyses to determine whether SV2C mediates rotenone-induced PD outcomes other than lethality.

Following the results of the lethality analysis, a sublethal dose of each toxicant equivalent to approximately 60% of the lowest dose tested in the lethality experiments was used to determine whether SV2C mediates toxicant-induced behavioral outcomes. Automated analysis of motor behavior was performed using WormLab software to determine the track length of individual worms across a 30s testing period. At 3.3mM MPP^+^, the GFP, GFP::α-syn, and GFP::SV2C strains showed no impairments in distance traveled compared to the solvent control (water) treated *C. elegans* of the same strain (Two-way ANOVA with Tukey’s post hoc multiple comparisons test, n.s.; Figure 4A). However, the GFP::α-syn::SV2C strain showed decreased distance traveled compared to the solvent control treated GFP::α-syn::SV2C *C. elegans* (p < 0.05) and compared to the MPP^+^ exposed GFP::α-syn (p < 0.05) (Two-way ANOVA with Tukey’s post hoc multiple comparisons test; Figure 4A).

**Figure 4.**
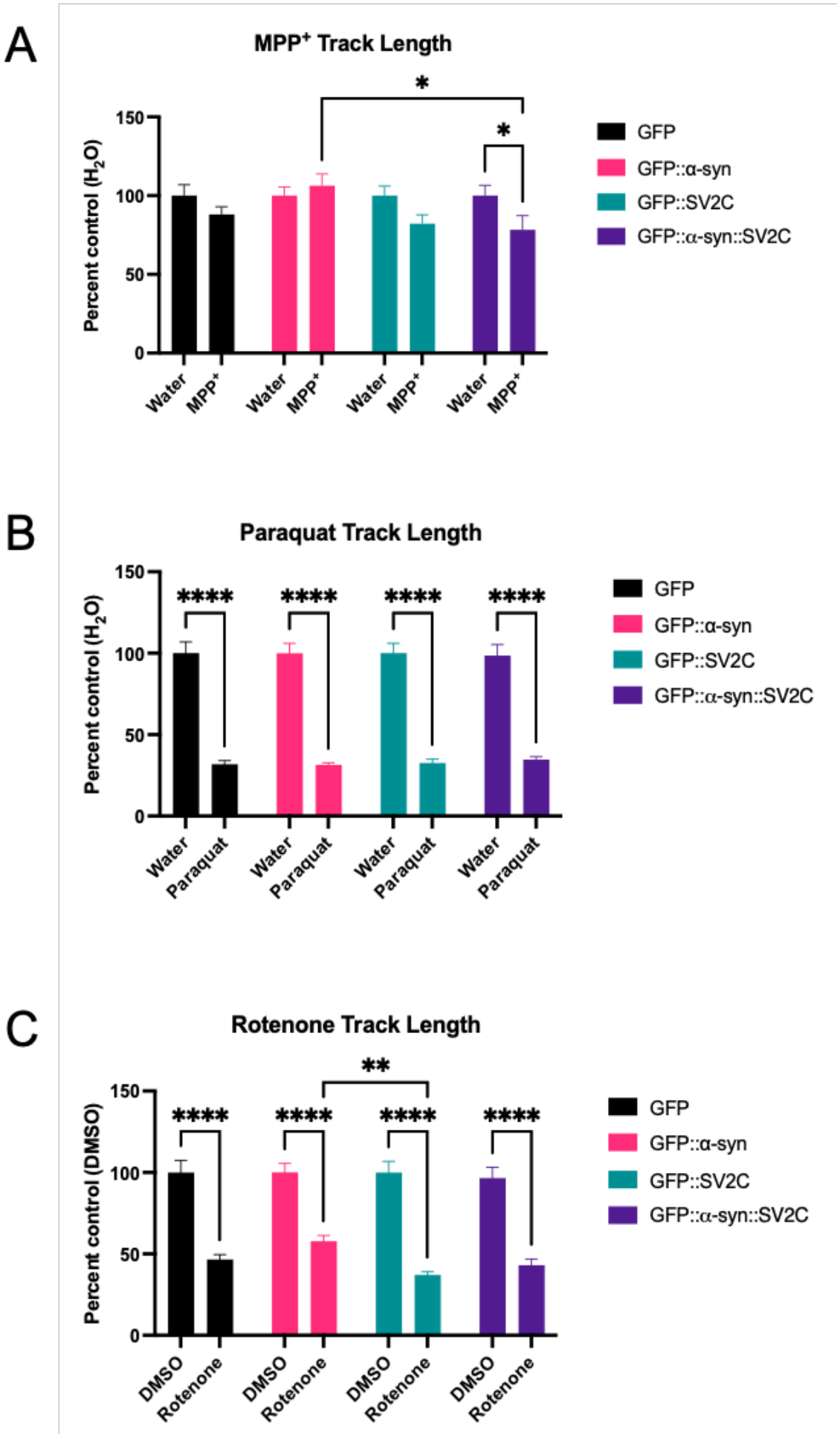
Behavioral effects following MPP^+^, paraquat, and rotenone exposure. A. Distance traveled in a 30s period following 48h exposure to 3.3mM MPP^+^ as represented by track length expressed as a percent of within strain solvent treated control. Data compiled across three experimental replicates with n = 55 - 82 *C. elegans* per group across three replicates. Data represented as mean and standard error of the mean. Two-way ANOVA with Tukey’s post hoc multiple comparisons test, *p < 0.05. B. Distance traveled in a 30s period following 48h exposure to 10mM paraquat as represented by track length expressed as a percent of within strain solvent treated control. Data compiled across three experimental replicates with n = 61 - 102 *C. elegans* per group across three replicates. Data represented as mean and standard error of the mean. Two-way ANOVA with Tukey’s post hoc multiple comparisons test, ****p < 0.0001. C. Distance traveled in a 30s period following 48h exposure to 13.3µM rotenone as represented by track length expressed as a percent of within strain solvent treated control. Data compiled across three experimental replicates with n = 56 - 90 *C. elegans* per group across three replicates. Data represented as mean and standard error of the mean. Two-way ANOVA with Tukey’s post hoc multiple comparisons test, **p < 0.01, ****p < 0.0001.

Following exposure to 10mM paraquat, all strains showed decreased distance traveled compared to the solvent control (water) treated *C. elegans* of the same strain (Two-way ANOVA with Tukey’s post hoc multiple comparisons test, p < 0.0001; Figure 4B); however, there were no differences when comparing between strains and treatments (Two-way ANOVA with Tukey’s post hoc multiple comparisons test, n.s.; Figure 4B). Similarly, at 13.3µM rotenone exposure, all strains showed decreased distance traveled compared to the solvent control (DMSO) treated *C. elegans* of the same strain (p < 0.0001) (Figure 4C). However, there was also a significant decrease in distance traveled between the GFP::SV2C strain and the GFP::α-syn::SV2C strain (Two-way ANOVA with Tukey’s post hoc multiple comparisons test, p < 0.01; Figure 4C).

These data suggest that despite increasing organism-level vulnerability to toxicant exposure, α-synuclein expression may confer protection against toxicant-induced behavioral deficits following sublethal exposure to MPP^+^ and rotenone. Likewise, although SV2C expression demonstrated organism-level protection to toxicant exposures, SV2C expression may exacerbate toxicant-induced behavioral deficits following sublethal exposure to MPP^+^ and rotenone. As motor behaviors such as distance traveled are mediated by dopamine transmission, and SV2C has been demonstrated to influence vesicular dopamine content, MPP^+^, paraquat, and rotenone were next evaluated for their impact on vesicular dopamine dynamics.

To determine whether SV2C expression impacted vesicular uptake activity, as previously observed with the VMAT2 inhibitor tetrabenazine, *in vitro* assays of vesicular uptake dynamics were performed.^11,38^ However, previous assays were performed in HEK cells expressing human VMAT2 and SV2C but without human dopamine transporter (DAT). Because MPP^+^ relies on DAT expression for cellular internalization, a new cell line was created expressing human DAT tagged with GFP, VMAT2, and SV2C (HEK-DAT-VMAT2-SV2C) for these experiments to be used in comparison to the parent cell line lacking SV2C expression (HEK-DAT-VMAT2) (Figure 5A).

**Figure 5.**
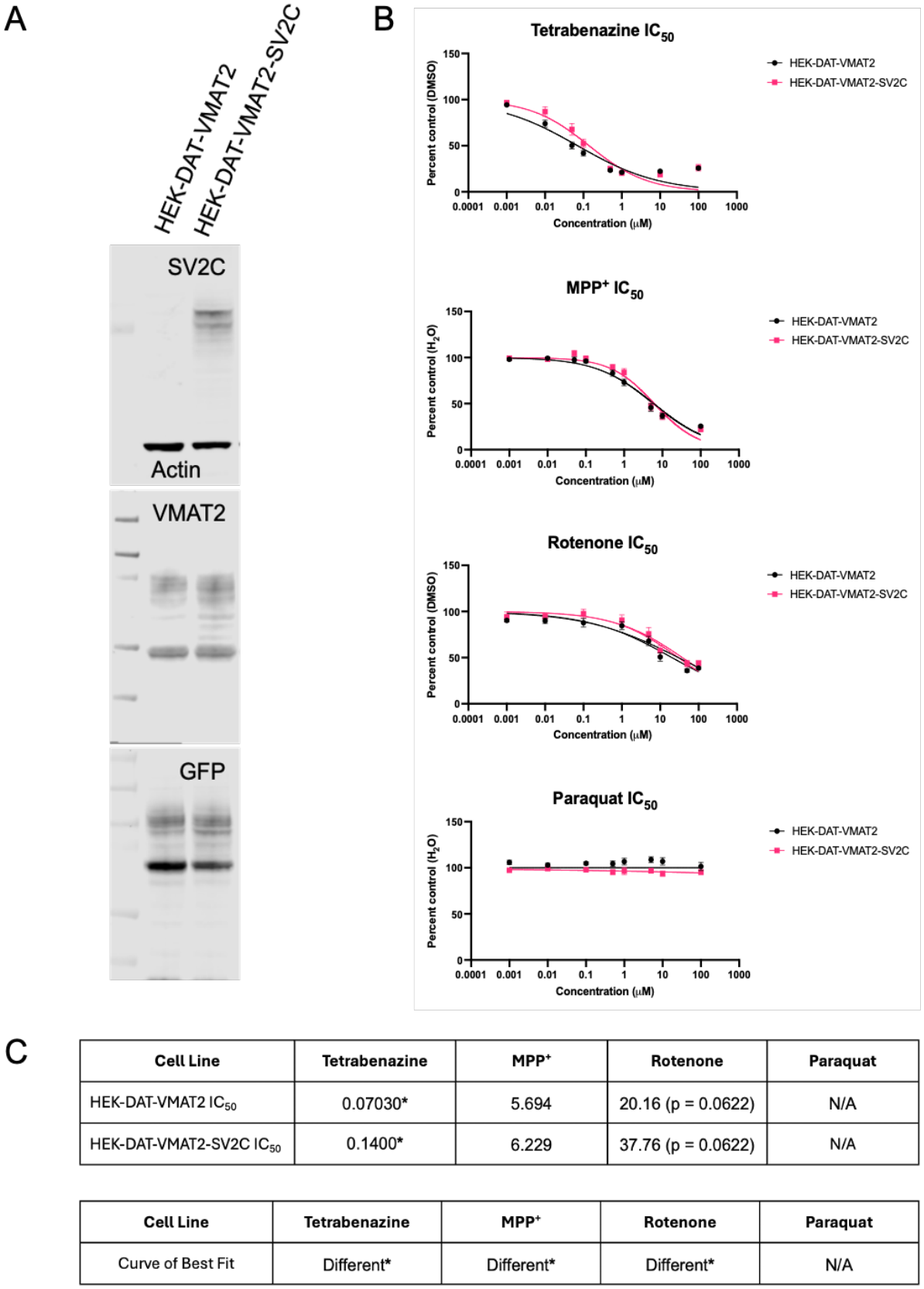
Effect of PD-associated toxicant exposures on vesicular uptake *in vitro*. A. Western blot showing SV2C, VMAT2, and GFP tag representing DAT expression in HEK-DAT-VMAT2 and HEK-DAT-VMAT2-SV2C cell lines. B. FFN206 uptake expressed as a percent on solvent control following dose-response exposure to tetrabenazine, MPP^+^, rotenone, and paraquat. For each experiment, there were 4–6 technical replicates for each cell and treatment condition, which were averaged to a single value for comparison across 4–8 experimental replicates. Nonlinear regression was performed to determine IC_50_ values and Curve of Best Fit. C. Results of nonlinear regression analysis IC_50_ calculation and Curve of Best Fit determination. Asterisks denote statistically significant differences. Tetrabenazine IC_50_ for HEK-DAT-VMAT2 = 0.0703 µM and HEK-DAT-VMAT2-SV2C = 0.140 µM (p < .05); Statistically Significantly Different Curve of Best Fit (nonlinear regression [inhibitor] versus normalized response, n = 4 experimental replicates for each dose, p < .05). MPP^+^ IC_50_ for HEK-DAT-VMAT2 = 5.694 µM and HEK-DAT-VMAT2-SV2C = 6.229 µM (p = n.s.); Statistically Significantly Different Curve of Best Fit (nonlinear regression [inhibitor] versus normalized response, n = 4-7 experimental replicates for each dose, p < .05). Rotenone IC_50_ for HEK-DAT-VMAT2 = 20.16 µM and HEK-DAT-VMAT2-SV2C = 37.76 µM (p = 0.0622); Statistically Significantly Different Curve of Best Fit (nonlinear regression [inhibitor] versus normalized response, n = 4 experimental replicates for each dose, p < .05). Paraquat IC_50_ for HEK-DAT-VMAT2 = N/A and HEK-DAT-VMAT2-SV2C = N/A (unable to calculate); Unstable curves for Curve of Best Fit (nonlinear regression [inhibitor] versus normalized response, n = 5 experimental replicates for each dose, N/A).

A dose response was performed for tetrabenazine, MPP^+^, rotenone, and paraquat and analyzed by nonlinear regression to determine the IC_50_ of each compound in the HEK-DAT-VMAT2 and HEK-DAT-VMAT2-SV2C cell lines (Figure 5B-C). Additional analysis was performed to determine if the Curve of Best Fit differed between the cell lines (Figure 5B-C). There was a significant difference in the IC_50_ for tetrabenazine, with HEK-DAT-VMAT2 IC_50_ = 0.0703 µM and HEK-DAT-VMAT2-SV2C IC_50_ = 0.140 µM, as well as a statistically significant difference in the curve of best fit for the two cell lines (nonlinear regression [inhibitor] versus normalized response, n = 4 experimental replicates for each dose, p < 0.05, Figure 5B-C). For MPP^+^, although there was no significant difference in the IC_50_ between the two cell lines, there was a statistically significant difference in the curve of best fit (nonlinear regression [inhibitor] versus normalized response, n = 4-7 experimental replicates for each dose, p < 0.05, Figure 5B-C).

Despite the rotenone lethality data in Figure 4 demonstrating inconsistent results, the *in vitro* assays demonstrated a trend toward a significant difference in the IC_50_ for rotenone, with HEK-DAT-VMAT2 IC_50_ = 20.16 µM and HEK-DAT-VMAT2-SV2C IC_50_ = 37.76 µM (p = 0.0622), and there was a statistically significant difference in the curve of best fit for the two cell lines (nonlinear regression [inhibitor] versus normalized response, n = 4 experimental replicates for each dose, p < 0.05, Figure 5B-C). Paraquat demonstrated no inhibition of uptake, preventing IC_50_ calculation and resulting in unstable curves for curve of best fit analysis (nonlinear regression [inhibitor] versus normalized response, n = 5 experimental replicates for each dose, N/A, Figure 5B-C).

## Discussion

In this study, we characterized novel *C. elegans* strains with transgenic expression of human SV2C and α-synuclein (Figure 1). Although SV2C is not endogenously expressed in *C. elegans*, they express a close homolog of SVOP, a protein related to the synaptic vesicular proteins (SV2).^39^ Like SV2, SVOP contains 12 transmembrane regions, however, they differ as SVOP is not highly glycosylated.^39^ While SVOP is a component of synaptic vesicles, its exact function is still undefined.^39^ The transgenic expression of SV2C within *C. elegans* is novel and provides a method to further investigate dopamine handling. We found that *C. elegans* expressing SV2C displayed a significantly shorter lifespan, with 0% alive by 22 days of adulthood compared to other strains. However, no overt morphological deficits, such as decreases in body length or impaired fecundity, were observed (Figure 2).

To determine whether SV2C protects against PD-outcomes in *C. elegans*, we investigated lethality and motor behavior following exposure to the PD-related toxicants MPP^+^, paraquat, and rotenone. SV2C expression showed moderate protection against organism-level lethality following MPP^+^ and paraquat exposure, with lethality exacerbated by the expression of α-synuclein (Figure 3). This is consistent with studies demonstrating that MPP^+^ induces α-synuclein oligomerization *in vitro*, and that α-synuclein knockout mice show resistance to MPTP toxicity.^3,40^ It is also consistent with findings that SV2C enhances vesicular storage of MPP^+^ *in vitro* and that loss of SV2C exacerbates 1-methyl-4-phenyl-1,2,3,6-tetrahydropyridine (MPTP)-induced toxicity in mice.^11^ As our previous research has demonstrated a positive effect of SV2C expression on the uptake of MPP^+^ within synaptic vesicles, the protection observed in *C. elegans* may be due to enhanced uptake of MPP^+^ in synaptic vesicles, which lessens mitochondrial complex I inhibition and prevents α-synuclein oligomerization, resulting in fewer neurotoxic effects and improved organism resilience.

SV2C expression is also protective against paraquat-induced lethality, particularly when α-synuclein is also expressed (Figure 3). Studies have suggested that paraquat increases the rate of α-synuclein aggregation *in vitro*, α-synuclein concentration in mice, and soluble misfolded α-synuclein aggregates in *Drosophila melanogaster*.^41–43^ This is consistent with our findings that the presence of α-synuclein significantly exacerbated lethality observed in paraquat-exposed *C. elegans*. Furthermore, SV2C expressing *C. elegans* showed the lowest lethality when exposed to paraquat compared to other strains. Aside from interaction with α-synuclein, paraquat is also thought to cause neurotoxicity through the generation of oxidative stress (as indicated by increased lipid peroxidation and reactive oxygen species (ROS) generation), and alterations to dopamine catabolism.^44^ Cytosolic dopamine is susceptible to enzymatic catabolism and oxidation, which also produce ROS.^25^ Building on findings that SV2C increases vesicular dopamine uptake and retention, SV2C may mitigate paraquat-induced toxicity by limiting cytosolic dopamine buildup, dopamine catabolism, and ROS generation.

Interestingly, the lethality observed following paraquat and MPP^+^ exposure in *C. elegans* was not replicated with rotenone (Figure 3). Rotenone also acts as a mitochondrial complex I inhibitor, contributing to reactive oxygen species (ROS) production, decreased levels of reduced glutathione antioxidant stores, and α-synuclein aggregation.^5^ Rotenone is highly lipophilic, allowing it to diffuse through biological membranes, which contributes to it being a significantly more potent complex I inhibitor than MPP^+^.^45,46^ Resistance to rotenone-induced toxicity in *C. elegans* has been observed previously, as demonstrated by the absence of significant dopaminergic neuron degeneration after 24 and 48 hours of exposure.^47^ Mitochondrial complexes II and V are upregulated in *C. elegans* following rotenone exposure and the glyoxylate pathway, an alternate variation to the TCA cycle in *C. elegans*, is also activated following 24 and 48 hour exposures to 1 μM and 5 μM rotenone, which are possible compensatory mechanisms preventing toxicity.^47^ While 48 hour exposure to rotenone does not cause significant toxicity, continuous exposure to 5 μM rotenone for 10 days causes mild dopaminergic neurodegeneration and increased vulnerability to ROS-producing metabolites.^48^ Thus, our findings of rotenone resilience in *C. elegans* are consistent with previous studies.

In *C. elegans*, analysis of the fluorescent marker for dopaminergic neurons can be used to evaluate neuron health through evaluation of neuron process integrity. Due to the pharynx marker in the transgenic SV2C strains, which overlaps with the dopaminergic neuron processes, we were unable to perform neuron health analyses to determine whether SV2C expression confers neuroprotection against PD-associated neurotoxicants. In this study, motor behavior as defined by the distance traveled in a 30s period was used as a proxy for dopaminergic neuron health (Figure 4). We found that exposure to paraquat and rotenone caused significant decreases in track length in all strains. However, track length deficits were not observed following MPP^+^ exposure.

The dose of exposures used in the behavioral assessments was chosen to observe sublethal effects of each compound. While previous work has identified motor deficits in *C. elegans* behavior following MPP^+^ treatment, these results followed a developmental exposure rather than exposure initiated in adulthood.^49^ Additionally, our previous work has demonstrated impairments in neuron health following sublethal exposure to MPP^+^.^50^ However, the deficit was only observed in the OK4111 (cat-1 null) strain, which lacks the ability to sequester MPP^+^ within vesicles and therefore have increased vulnerability to MPP^+^ exposure. Thus, the lack of behavioral effects observed with the dose of MPP^+^ used in this study suggests that a higher dose or a longer exposure duration may be required to observe MPP^+^-induced behavioral deficits, particularly to see differences based on strain genotype. On the other hand, the motor impairments observed following paraquat and rotenone exposure may have been too drastic to observe differences based on strain genotype, and future experiments with lower doses or shorter durations of exposure may provide insight into the ability of genotype to influence behavioral outcomes. Additionally, other measures of motor behavior such as movement speed or more nuanced motor features such as body bends, reversals, and turns may reveal toxicant-induced impairments and strain genotype-mediated differences.

The role of SV2C in mediating vesicular neurotransmitter uptake was evaluated *in vitro* using assays of fluorescent false neurotransmitter uptake (Figure 5). In a dose-response experiment, both MPP^+^ and rotenone caused inhibition of vesicular uptake. The inhibition of uptake due to toxicant exposure is similar to the inhibition of uptake observed with the VMAT2 inhibitor tetrabenazine, and as previously reported.^11,38^ However, no inhibition of uptake was observed with paraquat. Protective IC_50_ shifts were observed in cells expressing SV2C following MPP^+^ and rotenone exposure, suggesting that SV2C expression may protect against dysregulated dopamine sequestration caused by the competitive inhibitor MPP^+^ and non-competitive inhibitor rotenone. Despite not observing SV2C-mediated differences in behavior, this protection may explain the observed SV2C-mediated protection against toxicant-induced lethality.

MPP^+^ acts directly as a VMAT2 inhibitor by competing for uptake into synaptic vesicles with DA.^51^ Our previous work has demonstrated increased uptake of MPP^+^ in synaptic vesicles isolated from cells expressing human VMAT2 and SV2C, but this phenomenon has previously not been evaluated in intact cells.^11^ Previous studies have also demonstrated inhibition of VMAT2 activity by rotenone *in vitro*, and overexpression of *dVMAT* in *Drosophila melanogaster* preventing dopaminergic degeneration.^52–54^ The exact interaction between paraquat and VMAT2 remains unclear, however. While paraquat exposure reduces VMAT2 expression *in vitro*, overexpression of *dVMAT* does not protect against paraquat-induced dopaminergic cell death.^54,55^

This study characterizes novel *C. elegans* strains expressing human SV2C and α-synuclein and demonstrates the utility of these models for studying toxicant-induced PD outcomes in *C. elegans*. Our data supports existing evidence that implicates SV2C as a modifier of PD and lays the groundwork for future mechanistic studies to evaluate additional PD outcomes. Future experiments can test additional environmental exposures that have been linked to PD, and additional models of PD can be created by expressing SV2C in other *C. elegans* genetic models including LRRK2 expression. Mechanistic approaches, such as evaluating neuron health through non-fluorescent-mediated measures, α-synuclein aggregation, and metabolomic analyses can also provide further insight into the mechanisms of toxicity and SV2C-mediated protection.

## Acknowledgments

We thank Dr. Catharine Rankin for providing the UA44 strain and Dr. Jocelyn Dicent for advising experimental design in *C. elegans* experiments.

